# In vitro study of the growth, reproduction and pathogenicity responses of *Fusarium oxysporum* f. sp. *zingiberi* to autotoxins from ginger

**DOI:** 10.64898/2025.12.02.691968

**Authors:** Yan Zhang, Hongshen Guo, Yanping Xu, Xiaochuan Chen, Miaomiao Zhang, Naicheng Li, Abdullah Gera

**Author notes:** Corresponding author: Naicheng Li; Nc. li. wfit. edu.cn.

## Abstract

Long-term monoculture of ginger triggers *Fusarium* wilt, a disease caused by *Fusarium oxysporum* f. sp. *niveum* (*Foz*). However, the role of autotoxins in promoting pathogen growth remains unclear. Four representative autotoxins, syringic acid, coumarin, ferulic acid and 7-Hydrooxycoumarin were selected to investigate their allelopathic effects on the growth, reproduction, and virulence-associated traits of *Foz*. The responses of *Foz* to these compounds exhibited notable variations, which may be attributed to the structural differences among the autotoxins. The autotoxic compounds differentially enhanced the key pathogenic traits of *Foz*. Syringic acid was the most effective stimulant of mycelial growth and cell wall-degrading enzymes activity, concurrently boosting mycotoxin production via upregulating *FUB3* gene expression. Ferulic acid was the most potent promoter of sporulation and biomass accumulation, whereas 7-Hydroxycoumarin most effectively stimulated conidial germination. Notably, coumarin suppressed mycelial growth but strongly induced mycotoxin synthesis in *Foz*. These results provide mechanistic insights into how autotoxins in ginger monoculture systems promote *Fusarium* wilt.

## INTRODUCTION

Ginger (*Zingiber officinale* Roscoe) is an herbaceous plant that is generally cultivated as an annual crop despite its inherent perennial nature^[1]^. This ancient cultivar has been used for centuries in Asian cultures as a culinary spice and for therapeutic applications^[2]^. It exhibits anti-inflammatory, antitumor, antifungal, antibacterial, and anxiolytic properties, presenting significant economic opportunities for farmers^[3–5]^. The substantial economic benefits associated with ginger cultivation have incentivized the continuation of monoculture. Consequently, this has led to a progressive increase in the prevalence of pathogens such as *Fusarium* wilt^[6, 7].^

*Fusarium* wilt, caused by *Fusarium oxysporum* f. sp. *zingiberi* (*Foz*), was initially documented in Australia in the 1930s and is now prevalent in Asia, North America, and Oceania^[7, 8]^. The pathogen is categorized within the phylum Ascomycota and the genus *Fusarium*, and it causes vascular diseases in a variety of plant species, including melon, cucumber, and potato^[9–11]^. The infection is initiated when the fungus infiltrates the host’s vascular tissues through hyphal invasion, subsequently secreting enzymes that degrade the cell wall and producing mycotoxins^[6, 12]^. Research indicates that *Fusarium* spp. can persist saprophytically, enabling their survival in soil for prolonged durations in the absence of a host^[13]^. However, wilt manifests when the extent of root infection exceeds a critical threshold, thereby complicating control measures^[14]^. Concurrently, the activity and virulence of *Fusarium* spp. are also regulated by plant allelopathic compounds^[15, 16]^.

Allelopathy is a natural mechanism in plants that involves the release of chemicals, which affect the growth of other organisms^[17]^. Autotoxicity, a distinct form of intraspecific allelopathy, is characterized by the release of chemical compounds by plants that inhibit or suppress the germination and growth of conspecific individuals within the same species^[18]^. The allelochemicals released by the roots and decaying residues were isolated and identified, and were divided into different species based on their structures, including phenolic acids, aldehydes, coumarins, quinones, alkaloids, and terpenoids. Extensive research has been conducted on phenolic acids, which are believed to significantly contribute to the challenges encountered in continuous cropping systems^[19]^. These allelochemicals exert deleterious effects on seed germination, seedling development, root growth, plant quality, and resistance^[20, 21]^. Increasing evidence also suggests that autotoxicity may initiate and modulate pathogen occurrence, thereby mediating interactions and signaling processes between plants and microorganisms^[22, 23]^. For instance, cinnamic acids and coumarin exuded by watermelon roots have been documented to inhibit seedling growth while concurrently augmenting the pathogenicity factors of *Fusarium oxysporum* f. sp. *niveum*^[24, 25]^. These autotoxic chemicals have also been demonstrated to enhance mycelial growth, spore germination, and pathogenicity in *Fusarium oxysporum* f. sp. *melon*^[26]^. The application of exogenous vanillic acid and p-hydroxybenzoic acid facilitates the spore germination of *F. solani* and *F. equiseti*, while also modifying the structure of the rhizospheric soil microbial community^[27]^. These findings corroborate that plant root exudates facilitate wilt development.

Similar to other crops, ginger produces auto-toxic root exudates^[28]^. Although the phenomenon of auto-toxic root exudates is acknowledged, their precise role in the reproduction and virulence of pathogens remains unclear. Therefore, four autotoxic chemicals released by ginger, syringic acid, ferulic acid, coumarin and 7-Hydroxycoumarin, were explored and their mechanisms of action characterized. Changes in growth, reproduction and virulence factors of *Foz* were characterized.

## MATERIALS AND METHODS

### Pathogen strain and autotoxic compounds

The pathogenic strain JJF46 was isolated from infected ginger at the Laboratory of Plant Protection at the Weifang Institute of Technology, China. Morphological and molecular techniques were used to identify the pathogenic strain of *Fusarium oxysporum* f. sp. *zingiberi (Foz)*. Ferulic acid, coumarin, syringic acid, and 7-hydroxycoumarin have been used as autotoxic compounds, as documented in previous studies^[28]^. All chemicals were purchased from Aladdin Chemical Company (Shanghai, China). The doses and information of the deployed autotoxic chemicals are presented in Table 1.

**Table 1.** Dosage and information of autotoxic compounds used. Dimethyl sulfoxide (DMSO) was used as the solvent for all compounds.

| Autotoxic Chemicals | Dose (mmol L<sup>-1</sup>) | Purity | Structure |
| --- | --- | --- | --- |
| Ferulic acid | 0, (0.05), (0.1), 0.25, 0.5, (0.75), (1), (2) * | ≥99% | Phenolic acids |
| Coumarin | 0, (0.05), (0.1), 0.25, 0.5, (0.75), (1), (2) * | ≥99% | Lactone |
| Syringic acid | 0, (0.05), (0.1), 0.25, 0.5, (0.75), (1), (2) * | ≥98% | Phenolic acids |
| 7-Hydroxycoumarin | 0, (0.05), (0.1), 0.25, 0.5, (0.75), (1), (2) * | ≥99% | Lactone |

\*. The doses shown in parentheses were exclusively used in Experiment 1 or 2.

Two distinct concentration gradients of autotoxic compounds were established: the first was used to assess the allelopathic effects of these compounds on mycelial growth (Experiment 1), whereas the second was designed to evaluate their effects on sporulation, conidial germination, and pathogenicity-related factors based on the differential sensitivity of *Foz* to the autotoxic compounds (Experiment 2).

### Effects of autotoxic compounds on mycelial growth

To examine the impact of autotoxic compounds on mycelial growth, potato dextrose agar (PDA) was chosen as the medium for the administration of the autotoxic compounds. The final concentrations of these compounds in the PDA medium are listed in Table 1. A mycelial plug with a diameter of 6 mm, derived from a 7-day-old pure culture, was positioned at the center of each plate and incubated at 28 ℃ for 6 days. Mycelial growth was assessed using the cross method after 2, 4, and 6 d of incubation. The relative allelopathy intensity (RI) of mycelial growth was determined using Williamson’s method^[29]^. Each treatment was performed in five replicates.

### Effects of autotoxic compounds on sporulation

To assess the allelopathic impact of autotoxic compounds on sporulation, five agar plugs, each 6 mm in diameter, were extracted from a 7-day-old PDA medium and inoculated into Bilay and Joffe’s medium^[30]^. These were incubated at 28 ° C for 7 days in 250 ml flasks. The final concentrations of autotoxic chemicals present in the medium are listed in Table 1. After the 7-day incubation period, the broth was filtered through four layers of sterile gauze, and 1 mL of the filtrate was serially diluted to concentrations ranging from 10^-5^ to 10^-7^ ind ml^-1^. The resulting dilution was transferred to a hemocytometer for conidia enumeration using a microscope. Each treatment was conducted in quintuplicate.

### Effects of autotoxic compounds on conidial germination and biomass production

To investigate the impact of autotoxic compounds on conidial germination, *Foz* was cultured on potato dextrose agar (PDA) plates for seven days. Subsequently, five agar plugs, each 6 mm in diameter, were extracted from the culture and transferred to a liquid medium. The medium was incubated at 28 ° C for seven days under shaking conditions at 180 rpm. The resulting broth was filtered through sterile gauze and diluted with sterile water to achieve a concentration of ≤ 1000 conidia ml^-1^. A 100 μ L aliquot of the diluted suspension was spread onto PDA plates and incubated at 28 °C for two days. The number of colonies was counted to evaluate germination. Each treatment was performed in five replicates.

The filtered fungal mycelia were collected on sterile filter paper and desiccated at 80 ° C for 12 h until a constant mass was achieved. The corresponding filtrates were used for enzymatic assays. Each treatment was performed in five replicates.

### Effects of autotoxic compounds on cell wall-degrading enzymes

Pectinase, cellulase, protease, and amylase were selected as cell wall-degrading enzymes (CWDEs) of the pathogen in this study. Pectinase activity was assayed as described by Chang et al.^[31]^, with slight modifications. The reducing sugar released by enzyme hydrolysis of the polysaccharide substrate was reacted with 3,5-dinitrosalicylic acid (DNS) to produce a colored complex, which was quantified using an ultraviolet spectrophotometer. One unit of pectinase activity was defined as the amount of galacturonic acid (1 mg) produced by pectin breakdown per milliliter per hour. Cellulase activity was determined using the DNS method described by Brand and Alsanius^[32]^, where one unit of activity was defined as the amount of enzyme catalyzing the release of 1 μ g glucose per minute. Protease activity was measured following the method of Kole et al.^[33]^, with one unit defined as the amount of enzyme catalyzing the hydrolysis that produced 1 nmol of tyrosine per milliliter of sample per minute. Total amylase activity was assessed using the method described by Murado et al.^[34]^, where one unit of amylase activity was defined as the amount of enzyme that catalyzes the release of 1 mg of reducing sugar per mL of sample per minute.

### Effects of autotoxic compounds on mycotoxin

The response of mycotoxins to autotoxic compounds was assessed following the methodology outlined by Wu et al.^[24]^, with slight modifications. The experiment commenced with the establishment of a standard curve using fusaric acid (Sigma Chemical Co.) at specified concentrations. Five agar plugs, each 6 mm in diameter, were extracted from 7-day-old PDA plates and inoculated into Richard’s medium^[35]^, followed by incubation at 28 °C for 30 days. Post-cultivation, the broth was filtered through sterile gauze, and the pH was adjusted to 2 using 2 mol L^-1^ HCl. The medium underwent five extractions with equal volumes of ethyl acetate, allowing the contents to settle for 30 min after each extraction. The organic phase was centrifuged and decolorized using activated charcoal. Subsequently, the filtrate was dried and condensed at 28 ° C until a residue was obtained. The dried residue was reconstituted in 5 ml of ethyl acetate, and its optical density at 268 nm (OD_268_) was measured using ultraviolet spectrophotometry.

### Effects of autotoxic compounds on the expression of mycotoxin biosynthetic genes

To elucidate the allelopathic effects of autotoxic compounds on mycotoxin biosynthesis, the *FUB* gene cluster was selected for detailed examination and characterization ^[36]^. This study included five genes from this cluster: *FUB1*, *FUB3*, *FUB6*, *FUB8*, and *FUB9*. The primer sequences and functional descriptions of these genes are listed in Table 2. A 6-mm-diameter agar plug obtained from a 7-day-old PDA culture was inoculated onto a fresh PDA plate supplemented with the corresponding autotoxic chemicals and incubated at 28 °C for 7 days. Subsequently, mycelia were harvested for gene expression analysis.

**Table 2.**
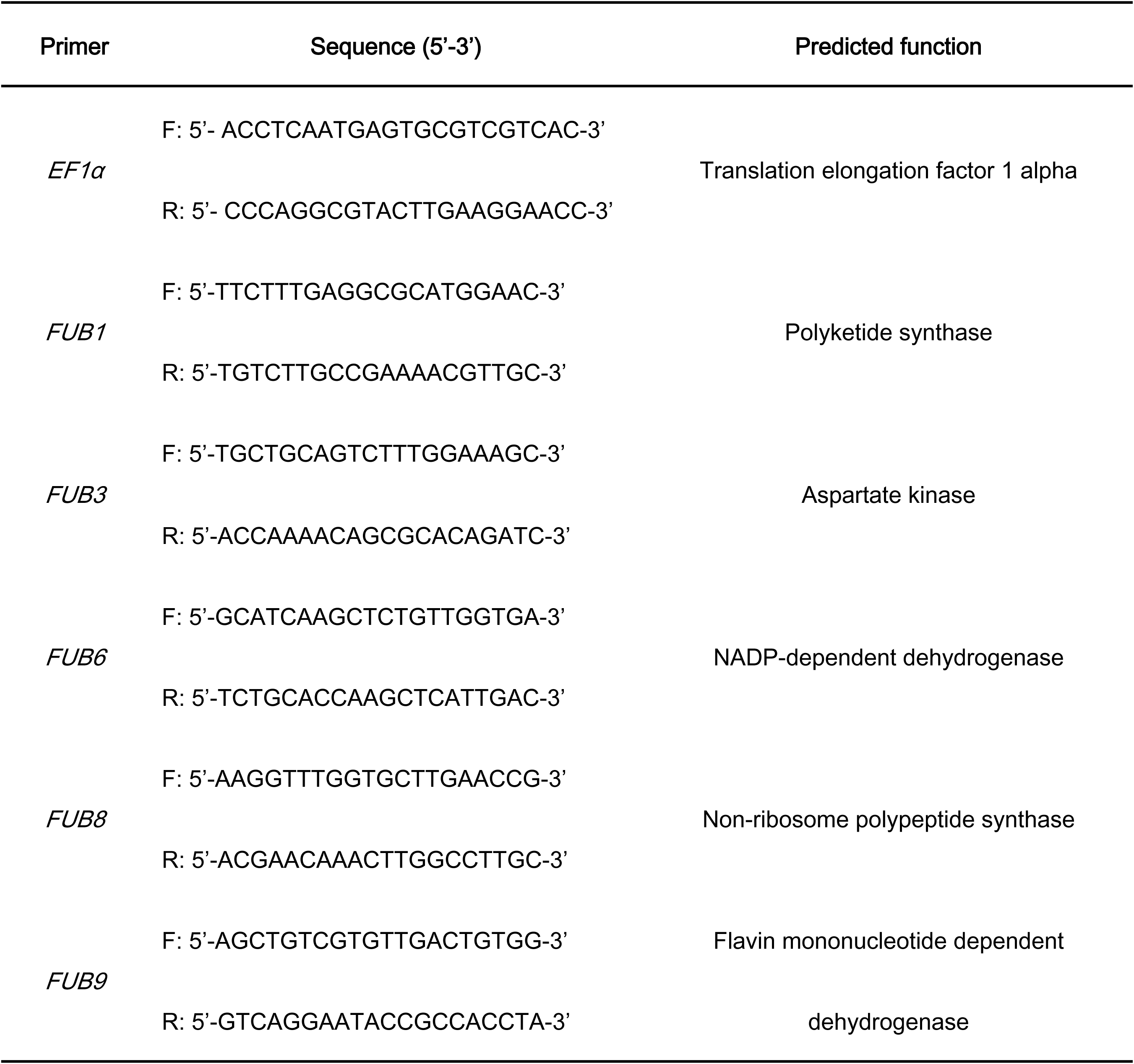
Primer pair Sequences and corresponding gene functions in the pathogen used for real-time PCR analysis.

Total RNA was extracted using a commercial MiniBEST Plant Total RNA Extraction Kit (TaKaRa, Dalian, China), verified by electrophoresis, and quantified using a NanoDrop spectrophotometer (Thermo Fisher, Germany). First-strand cDNA synthesis was performed using the Prime Script™ II 1st stand cDNA Synthesis kit (TaKaRa, Dalian, China). Quantitative real-time PCR (qRT-PCR) was performed using the TB Green™ Fast qPCR Mix kit (TaKaRa, Dalian, China) according to the manufacturer’s instructions, as described by Li et al.^[37]^. Translation elongation factor 1 alpha (*EF1α*) was used as the internal reference^[36]^. Relative gene expression levels were calculated using the 2^−△△Ct^ method.

### Statistical analysis

The allelopathic effects of autotoxic substances on fungal development, including mycelial growth, sporulation, colonial germination, and biomass production, were quantified using the relative intensity index (RI). The RI index was determined using the following equations: RI = 1 − C/T (T≥C) and RI = T/C − 1 (T<C), where C and T denote the control and treatment data, respectively. According to this index, a positive RI value (RI >0) signifies stimulation, whereas a negative RI value (RI <0) indicates inhibition^[29]^. To assess the allelopathic effects of autotoxic substances on cell wall-degrading enzymes (CWDEs), a weighted approach was employed for four enzymes based on their relative significance in the infection process^[38]^. The weights assigned to cellulase and pectinase were 0.3 each, while those assigned to protease and amylase were 0.2 each. The allelopathic effects of autotoxic chemicals on CWDEs were quantified using the equation: PI = T/C × W, where C and T represent the control and treatment data, respectively. W denotes the weight, and PI represents the allelopathic intensity index. To comprehensively evaluate the allelopathic effects of autotoxic chemicals, the cumulative RI and PI indices were calculated as integrated measures.

A multiple comparison tests were performed using SPSS 22.0 software to assess significance between the control and treatment groups The significance level (P) were set at *P* < 0.05 and *P* < 0.01.

## RESULTS

### Effect on mycelial growth and biomass production

The hyphal growth response exhibited variability among the autotoxic chemicals (Fig. 1a-d). Mycelial growth remained stable, with a mean of 2.63±0.34 (n=65), under low-dose treatments (≤ 0.25 mmol L^-1^) for all chemicals over a 2-day incubation period, except for syringic acid, which significantly enhanced fungal growth at concentrations ≤ 0.1 mmol L^-1^. However, a marked inhibitory effect was observed at a higher concentration (0.5 mmol L^-1^) of syringic acid. After 4 days of incubation, colony growth generally increased relative to the controls under low-dose treatments but was suppressed at higher concentrations, particularly with syringic acid, where the colony diameter was reduced by 30.36% at 0.5 mmol L^-1^ compared to the controls. Consistent with the previous trend, fungal growth was significantly stimulated by low-dose treatments of all autotoxic chemicals except 7-Hydroxycoumarin over a 6-day incubation period. The most pronounced stimulatory effect was observed with syringic acid at 0.05 mmol L^-1^, where the colony diameter reached 7.83 cm after 6 d of incubation. In contrast, the high-dose treatment inhibited mycelial growth, corroborating the results observed after 4 days of exposure. At a coumarin concentration of 0.5 mmol L^-1^, the colony diameter was reduced to 4.82 cm compared to 6.38 cm in the control.

**Fig. 1.**
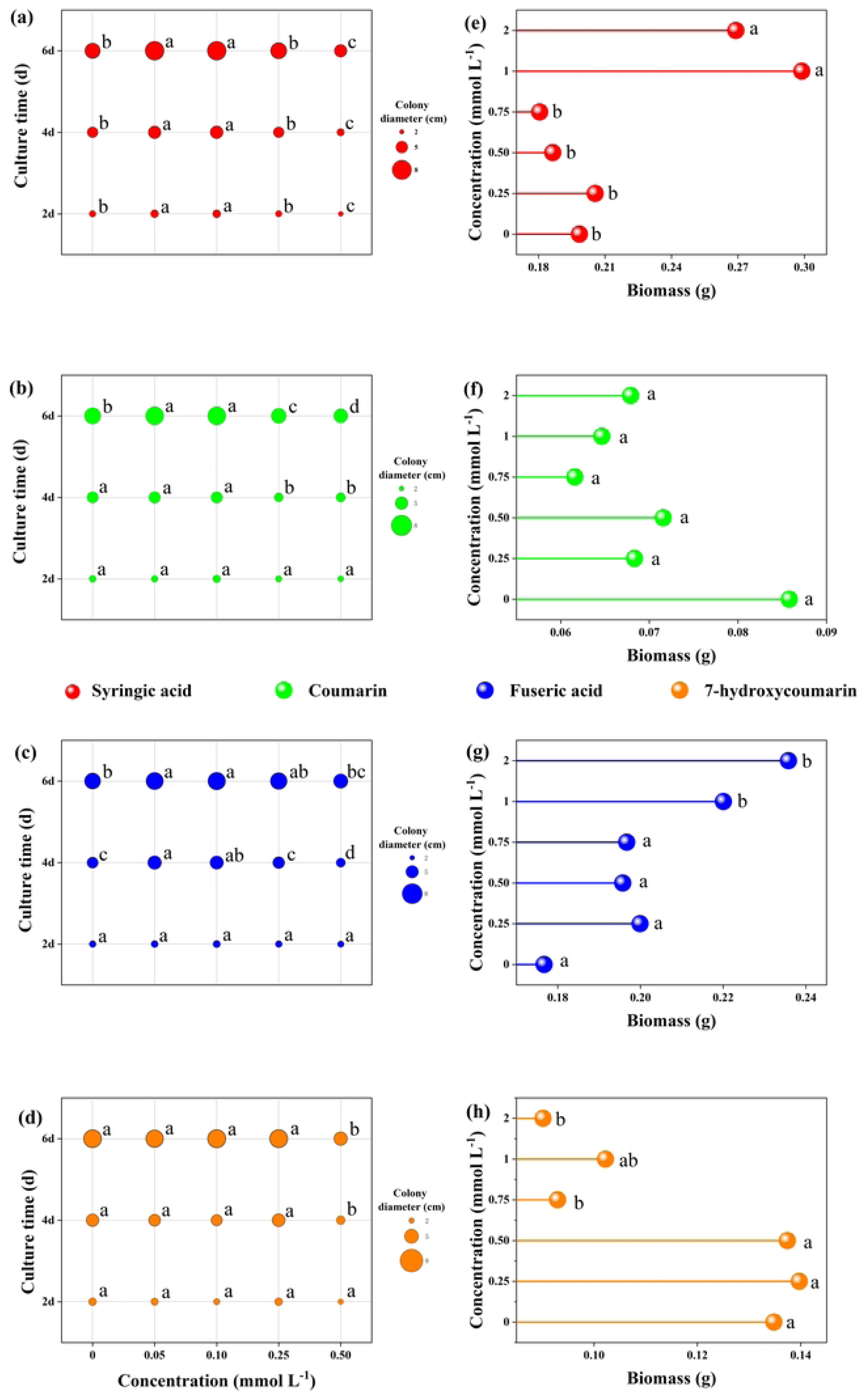
Impact of various autotoxins on colony growth (a-d) and biomass production (e-h) of *Fusarium oxysporum* f. sp. *niveum* (*Foz*). The autotoxins tested were (a, e): Syringic acid; (b, f): Coumarin, (c, g): Ferulic acid, and (d, h): 7-Hydroxyccoumarin. Different lowercase letters above bars indicate significant differences(*P*<0.05).

Biomass production did not exhibit the same pattern as mycelial growth (Fig. 1e-h). Coumarin had no significant impact on biomass accumulation, with the dry weight of the mycelia remaining constant at 0.07 ± 0.01 g. At lower concentrations of the other chemicals, no significant changes were observed after 7 days of incubation; however, distinct responses were observed at higher concentrations. Ferulic and syringic acids significantly increased fungal biomass compared to the controls by 33.45% and 35.59%, respectively, at concentrations ≥1 mmol L^-1^. Conversely, 7-Hydroxycoumarin demonstrated an inhibitory effect, as evidenced by a marked reduction in biomass at concentrations ≥0.75 mmol L^-1^.

### Effect on sporulation and spore germination

Sporulation responses exhibited significant variation across the different chemical treatments (Fig. 2a-d). At low concentrations ( ≤ 0.75 mmol L^-1^), coumarin had minimal influence on sporulation. However, with increasing coumarin concentrations, spore production was markedly enhanced, increasing from 38.67× 10⁶ in the control to 90× 10⁶ at 2 mmol L^-1^. A similar positive effect was observed with ferulic acid, where sporulation increased from 44.33× 10⁶ in the control to 72.67× 10⁶ at 1 mmol L^-1^, although this stimulation diminished at higher concentrations. The effects of syringic acid and 7-Hydroxycoumarin followed a comparable pattern: sporulation was significantly promoted at low concentrations ≤ 0.75 mmol L^-1^, but declined at higher concentrations (2 mmol L^-1^). At the highest concentration, spore numbers in the liquid medium decreased to 7.37 × 10⁶ and 3.73 × 10⁶ for syringic acid and 7-Hydroxycoumarin, respectively. Ferulic acid (Fig. 2g) did not induce significant changes in spore germination, with colony numbers remaining relatively stable at 83.97±2.67× 10⁶ (n=30). A slight increase was observed in the coumarin treatment at 0.75 and 1 mmol L^-1^ (Fig. 2f). In contrast, syringic acid (Fig. 2e) and 7-Hydroxycoumarin (Fig. 2h) significantly enhanced spore germination at low concentrations, with 7-Hydroxycoumarin demonstrating a clear concentration-dependent response. However, both compounds exhibited mild inhibitory effects at their highest concentrations. Specifically, spore germination under syringic acid declined from 74.50× 10⁶ in the control to 62.25× 10⁶ at 2 mmol L^-1^, while under 7-Hydroxycoumarin, it decreased from 99.00× 10⁶ in the control to 73.50× 10⁶ at 2 mmol L^-1^.

**Fig.2.**
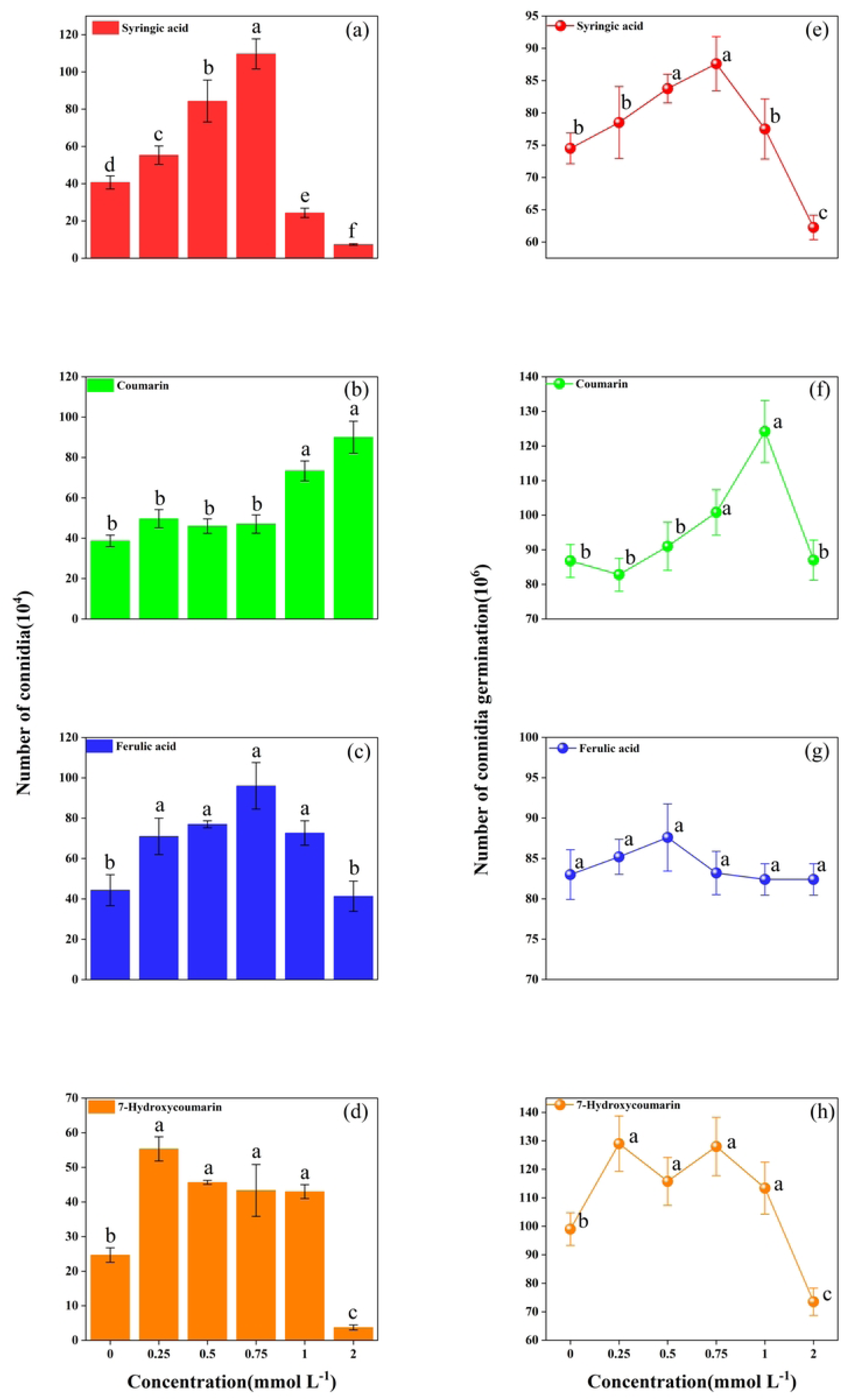
Effects of different autotoxins concentrations on sporulation and conidial germination of *Fusarium oxysporum f. sp. niveum (Foz).* Effects on sporulation by (a): Syringic acid (b): Coumarin (c): Ferulic acid and (d): 7-Hydroxyccoumarin. Effects on conidial germination by (e): Syringic acid (f): Coumarin (g): Ferulic acid (h): 7-Hydroxyccoumarin. Different lowercase letters above bars indicate significant differences (*P*<0.05).

### Impact on the activity of cell-wall degrading enzymes

After 7 days of exposure to varying concentrations of autotoxic compounds, pectinase activity showed distinct responses to each treatment (Fig. 3a). In the presence of ferulic acid, enzyme activity was significantly inhibited compared with the control by 65.89%, 52.64%, 49.52%, 75.61%, and 30.50% at concentrations of 0.25–2 mmol L^-1^, showing strong, concentration-dependent inhibition. In contrast, both coumarin and syringic acid markedly stimulated pectinase activity at concentrations ≥0.75 mmol L^-1^, with syringic acid having the most pronounced effect. This suggests an enhanced infection potential of the pathogen due to increased enzyme production, even at a low concentration (0.5 mmol L^-1^) of the elicitor. For 7-Hydroxycoumarin, pectinase activity initially increased from 0.45 mg mL⁻¹ h⁻¹ at 0.25 mmol L⁻¹ to 0.57 mg mL⁻¹ h⁻¹ at 0.75 mmol L⁻¹, but subsequently declined with higher doses, reaching a minimum of 0.21 mg mL⁻¹ h⁻¹ at the highest concentration.

**Fig. 3.**
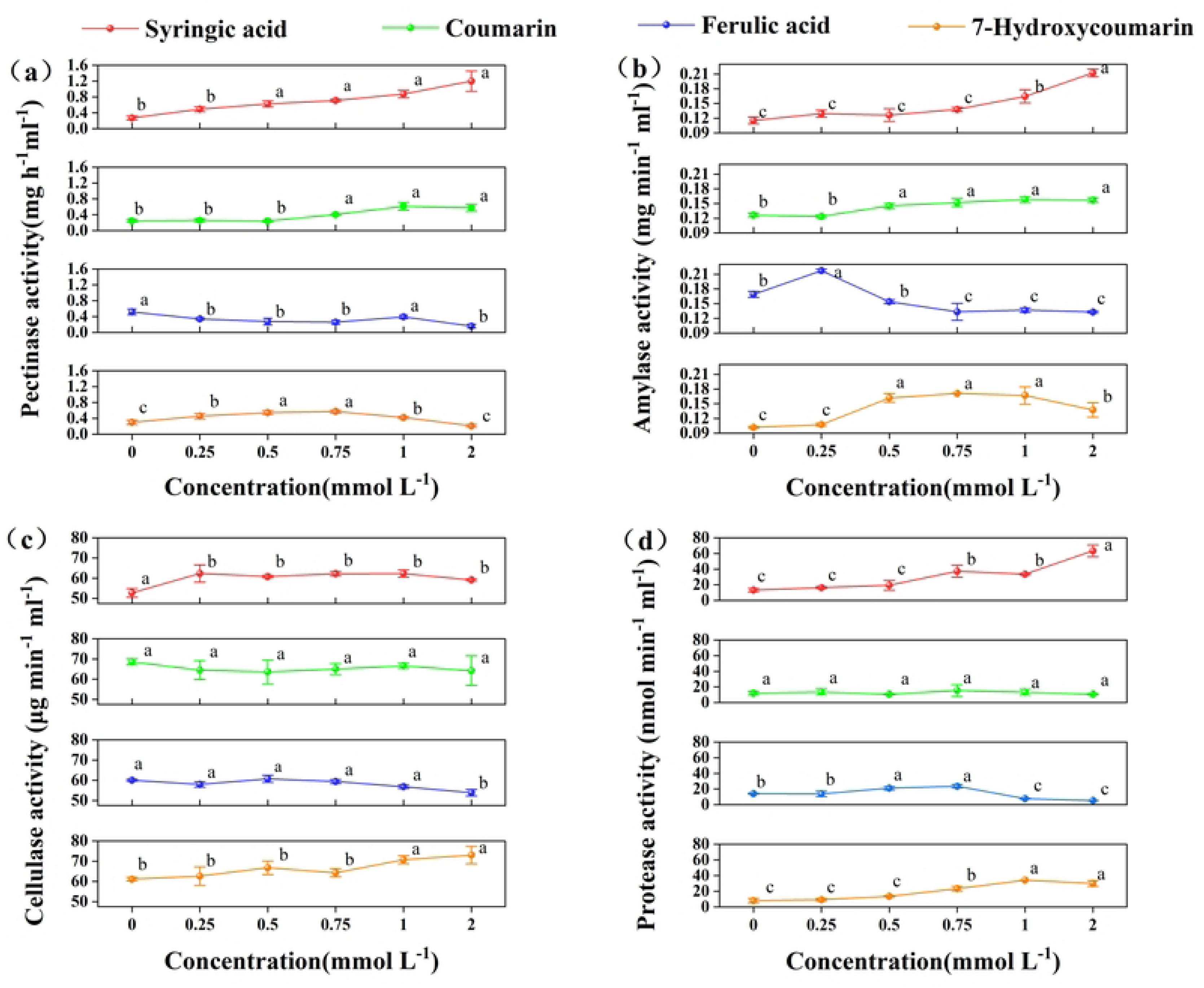
Effects of autotoxins at different concentrations on activity of cell-wall degrading enzymes. (a): Pectinase activity; (b): Amylase activity; (c): Cellulase activity; (d): Protease activity. Different lowercase letters indicate significant differences at (*P*<0.05).

In the presence of syringic acid, amylase activity was significantly stimulated in a concentration-dependent manner, increasing from 0.12 to 0.21 mg mL⁻¹ min⁻¹ across all treatment levels (Fig. 2b). Similar positive effects were observed under 7-Hydroxycoumarin and coumarin treatments at concentrations ranging from 0.5 to 2 mmol L^-1^, although a slight decline was detected at the highest concentration of 7-Hydroxycoumarin. In contrast, ferulic acid exhibited a dose-dependent effect, significantly enhancing amylase activity at the lowest concentration and inhibiting it at higher concentrations.

Syringic acid significantly induced cellulase activity (Fig. 3c), a key enzyme in pathogen infection, although not in a concentration-dependent manner. Activity remained consistently elevated, averaging 61.35 μg^-1^ ml^-1^ min^-1^ (n=15). Ferulic acid and 7-Hydroxycoumarin had minimal effects, with only a slight increase in activity observed at concentrations ≥2 mmol L^-1^ and ≥1 mmol L^-1^, respectively.

Analogous to other enzymatic responses, protease activity in syringic acid treatments (≥0.75 mmol L^-1^) increased significantly from 13.12 nmol mL⁻¹ min⁻¹ in the control treatment to 63.32 nmol mL⁻¹ min⁻¹ at 2 mmol L^-1^ (Fig. 3d). Similarly, protease activity was markedly enhanced in coumarin treatments (≥ 0.75 mmol L^-1^), showing a 377.66% increase at 2 mmol L^-1^. In contrast, ferulic acid promoted protease activity at low concentrations (≤ 0.75 mmol L⁻¹) but significantly inhibited it at higher concentrations (37.86% at 2 mmol L^-1^). No significant changes were detected in coumarin treatments, and enzyme activity remained within the range of 12.49±3.04 nmol mL⁻¹ min⁻¹.

### Effect on mycotoxin production

Mycotoxin production was strongly stimulated by most autotoxins in liquid culture, with the exception of 7-Hydroxycoumarin which exerted an inhibitory effect (Fig. 4). The stimulatory effects of syringic acid and coumarin were found to be concentration-dependent. At their highest concentrations, syringic acid and coumarin yielded highest concentration syringic acid and coumarin resulted in mycotoxin levels of 36.98 μg ml^-1^ and 38.56 μg ml^-1^, corresponding to 2.99 and 3.28 fold increases over the control (CK), respectively. A similar promotive effect was observed for ferulic acid, which increased mycotoxin production from 10.90 μ g ml^-1^ to 27.19 μ g ml^-1^. Conversely, 7-Hydroxycoumarin progressively reduced mycotoxin yields with increasing concentration.

**Fig. 4.**
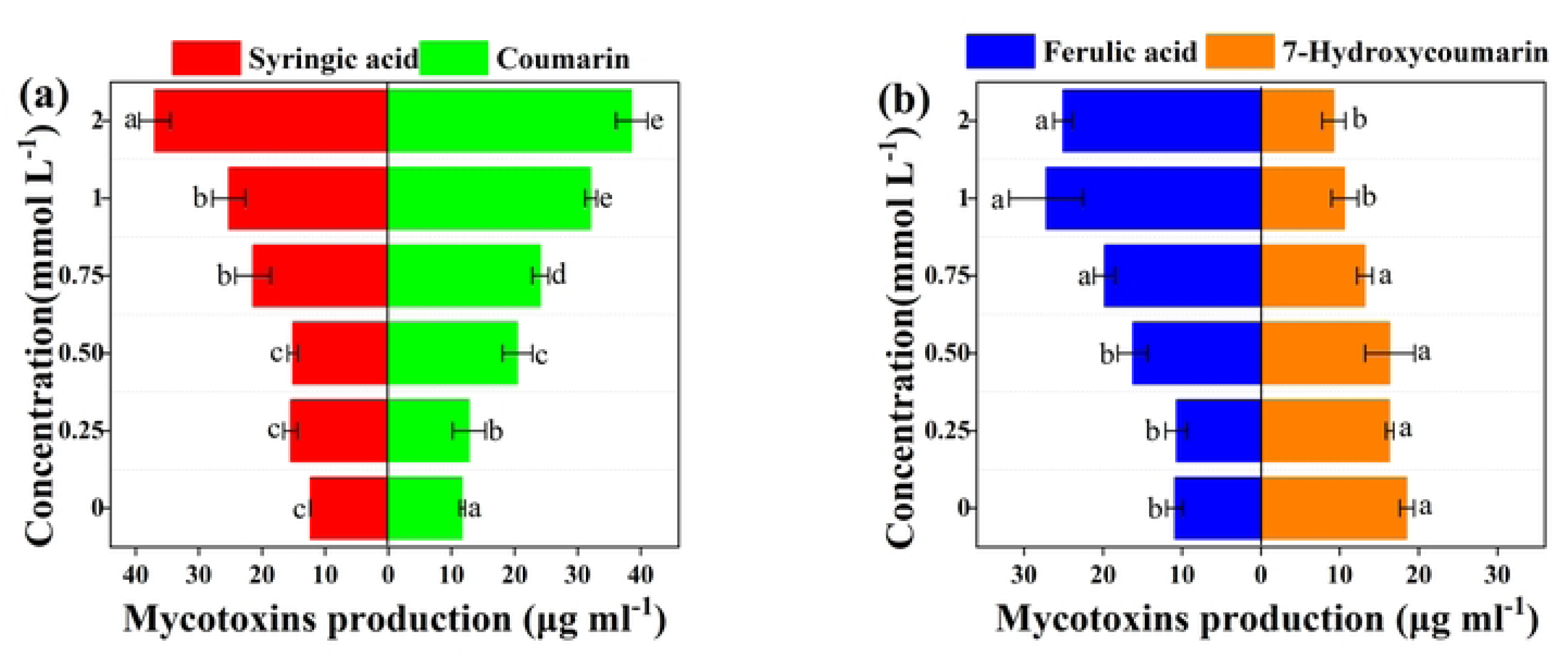
Mycotoxin production in response to different concentrations of autotoxins. (a): Syringic acid and Coumarin (b): Ferulic acid and 7-Hydroxycoumarin Different lowercase letters indicate significant differences among treatments (*P*<0.05).

### Impact on the expression of mycotoxin biosynthetic genes

The expression of mycotoxin biosynthetic genes varied under different autotoxin treatments. Syringic acid markedly induced mycotoxin accumulation, as evidenced by the gradual upregulation of *FUB3* and *FUB9*, which increased by approximately twofold relative to the control. In contrast, the relative expression levels of *FUB1*, *FUB6* and *FUB8*, showed only slight increases in response to syringic acid. Coumarin strongly upregulated *the FUB* gene cluster, particularly *FUB3*, *FUB6*, and *FUB9*, at the highest treatment concentration (2 mmol L^-1^), showing 2.41-, 2.64-, and 2.90-fold increases compared to the control, respectively. *FUB1* and *FUB8* expression levels also increased at coumarin concentrations ≥ 1 mmol L^-1^. In the presence of ferulic acid, the relative expression of *FUB9* initially increased significantly at lower concentrations (≤1 mmol L^-1^) and then declined to 2.43-fold of the control at the highest concentration. At 0.75 mmol L^-1^ and 1 mmol L^-1^, *FUB9* expression reached 5.43-and 6.31-fold of the control, respectively. A similar trend was observed *for FUB6* and *FUB8*. The *FUB3* gene displayed a comparable response, although the stimulatory effect was weakened at 1 mmol L^-1^. In contrast, 7-Hydroxycourmarin treatment led to an overall downregulation of FUB genes. There was a reduction in the expression of *FUB6* declined from 0.96 at 0.25 mmol L^-1^ to 0.77 at 2 mmol L^-1^ while *FUB9* expression decreased sharply, reaching its lowest level (0.52) at 0.75 mmol L^-1^. *FUB3* and *FUB8* expression remained largely unchanged, and *FUB1* showed only slight induction in response to 7-Hydroxycourmarin.

### Comprehensive effect of different autotoxins on *Fusarium oxysporum* f. sp. *niveum* (*Foz*)

The allelopathic effect of autotoxins on *Foz* varied distinctly (Fig. 6). Syringic acid was the most comprehensive stimulant, strongly enhancing all measured aspects of pathogenicity, particularly mycelial growth and CWDEs. The RI indices were 0.24 and 1.88, respectively. Coumarin strongly induced mycotoxin synthesis but concurrently reduced mycelial growth and biomass and had a slight inhibitory effect on CWDEs. Ferulic acid significantly increased sporulation and biomass but had a slight inhibitory effect on CWDEs. Conversely, 7-Hydroxycoumarin had the most divergent effects; it was the strongest enhancer of conidial germination but also the strongest inhibitor of mycelial growth, biomass accumulation, and mycotoxin production.

## DISCUSSION

Autotoxicity, a prevalent natural phenomenon, is recognized as a significant factor contributing to soil sickness in numerous crops^[39]^. Various autotoxic compounds have been identified that exert substantial inhibitory effects on plant growth and yield, primarily by exacerbating soil-borne diseases. Phenolic acids, coumarins, and esters have been identified as the principal autotoxins^[40–42]^. In the context of ginger, previous research has demonstrated that ginger root exudates markedly suppress seedling growth, impair antioxidant enzyme function, and alter membrane permeability^[28]^. The present study elucidated the allelopathic effects of autotoxins on pathogenic fungi. These findings indicate that the four chemicals exhibited distinct effects on *Fusarium* wilt in ginger, likely attributable to variations in their molecular structures. At specific concentrations, these compounds promoted mycelial growth, spore germination, cell wall-degrading enzyme activity, and mycotoxin production.

*Fusarium oxysporum* f. sp. *zingiberi* (*Foz*) is a common soil-borne pathogen that causes ginger wilt, a disease characterized by leaf withering and plant wilting^[8]^. The molecular mechanisms underlying *Foz* infection have been widely studied^[43]^. Under natural conditions, this soil-borne disease typically progresses through several phases. The initial phase is often triggered by root exudates from the host plant, such as phenolic and amino acids, which signal the pathogen to initiate its invasion program by stimulating spore germination and germ tube growth^[44]^. Second, the germ tube of the pathogen colonizes the root surface, establishing the first physical contact^[45]^. It then attaches to and forms a mycelial network that proliferates throughout the host root system. In the third phase, the fungus secretes CWDEs to break down the host’s defenses and invade the root cortex and vascular tissues, including the xylem vessels^[46]^. Finally, pathogens release toxins and other virulence factors to induce disease symptoms in the host plant^[12]^. Therefore, to elucidate the role of autotoxic chemicals in promoting plant disease incidents, it is essential to first interpret the allelopathic effects of these chemicals on the physiological activities of pathogens.

In a natural environment, the growth and reproduction of plant pathogens within the host represent their fitness and survival capacity^[47]^. These physiological processes are regulated not only by the pathogens themselves but also by root exudates secreted by the host plants. In the present study, most autotoxic chemicals followed a typical hormetic response, except 7-Hyoxycoumarin, exhibited similar effects on mycelial growth, where low concentrations (<0.25 mmol L^-1^) promoted and high concentrations ( ≥ 0.25 mmol L^-1^) inhibited mycelial growth (Fig.1), consistent with previous reports^[48]^. However, this stimulatory effect of the other phenolic acids diminished over time, as indicated by the reduction in the mycelial growth rate on solid medium. This phenomenon may be attributed to the fact that gradually increasing concentrations could suffice to inhibit pathogen growth. Previous research has shown that numerous plants counteract potential soil-borne pathogens by altering the quantity and composition of organic acids in their root exudates^[49–51]^. Therefore, the diminished promotive effect of these chemicals may represent one of the mechanisms through which ginger defends against this pathogen. Furthermore, the effect on biomass was inconsistent with that on mycelial growth. Ferulic and syringic acids increased biomass accumulation at high concentrations (> 1 mmol L^-1^), coumarin and 7-Hydroxycoumarin significantly inhibited it (Fig.1). This discrepancy is likely due to the structural differences among the chemicals. Specifically, phenolic acids (i.e. ferulic acid and syringic acid) may have acted as additional carbohydrate sources for the pathogen, thereby stimulating biomass production^[52]^.

Spores are essential reproductive units in fungi^[53]^. Previous research has established a correlation between spore content and the pathogenicity of *Fusarium* spp.^[12]^. In the present study, autotoxic chemicals exhibited a consistent pattern of sporulation and colonial germination, characterized by stimulation at low concentrations and inhibition at high concentrations, with the exception of ferulic acid (Fig. 2). This phenomenon may explain the increased incidence of Fusarium wilt in ginger cultivated in monoculture systems^[22]^. Concurrently, this promotion may be driven by the pathogen’s capacity to remodel host metabolism and induce the production of metabolites that facilitate pathogen reproduction^[54]^. Moreover, high coumarin concentrations notably stimulated sporulation and conidial germination (Fig. 2b and f). This response may be attributed to the inhibition of the deacetylase FoSir5 of *F. oxysporum* induced by coumarin. A decline in FoSir5 activity increases ATP production, enabling dormancy break and initiating germ tube formation^[55]^.

Cell wall-degrading enzymes (CWDEs) are critical virulence factors in plant-pathogenic fungi^[38]^. In *Fusarium oxysporum*, the enzymatic activities of pectinases and cellulases are integral to the degradation of pectin and cellulose, thereby facilitating host invasion^[56, 57]^. Previous research has also identified amylases and proteases as pathogenicity factors, with gene knockouts resulting in the total or partial loss of pathogenicity in *F. oxysporum*^[58, 59]^. In the present study, syringic acid exhibited the most pronounced stimulatory effect on these hydrolytic enzymes, even at low concentrations (≥0.25 mmol L-1) (Fig. 3). Both pectinase and proteinase activities were significantly enhanced by 7-Hydroxycoumarin across all treatments, potentially facilitating *F. oxysporum* infection and dissemination within host tissues. The increased activity of these enzymes aids *F. oxysporum* in hydrolyzing polymers and compromising the integrity of plant cell wall polymer. Consequently, the stimulatory effect of autotoxic chemicals on hydrolytic enzyme activity is considered a mechanism underlying the enhanced pathogenicity of *F. oxysporum*. Additionally, coumarin and ferulic acid also stimulated pectinase and amylase activities at specific concentrations. However, this effect is complex, as ferulic acid strongly inhibited pectinase activity across all treatments, corroborating previous findings^[60]^. This apparent contradiction underscores the context-dependent influence of phenolic compounds, which can either promote or suppress the secretion, synthesis, and activity of CWDEs.

The secretion of secondary metabolites by fungal plant pathogens is essential for their survival and adaptation to dynamic environments. *Fusarium* spp., which are model plant pathogens, can produce a range of mycotoxins that act as critical virulence factors or effectors, leading to plant wilting^[61]^. Among these, fusaric acid (FA), also known as 5-butylpicolinic acid, is a polyketide-derived secondary metabolite commonly synthesized by *Fusarium* species and is closely associated with wilt symptoms in host plants^[43]^. The phytotoxicity of FA is dependent on its concentration; moderate doses significantly inhibit plant growth and reduce biomass, whereas high concentrations result in severe leaf wilting and necrosis^[42]^. The biosynthesis of fusaric acid is regulated by a gene cluster consisting of 12 genes, collectively referred to as the *FUB* cluster, whose expression directly affects FA production^[62]^. Previous studies have identified *FUB1*, *FUB3*, and *FUB6* as essential genes involved in FA synthesis^[36]^. In the present study, the majority of autotoxic compounds demonstrated a pronounced stimulatory effect on FA in *Fusarium oxysporum* f. sp. *zingiberi* (*Foz*), with the exception of 7-Hydroxycoumarin (Fig. 4). However, the mechanisms underlying stimulation by autotoxic compounds appear to vary, as evidenced by distinct gene expression patterns (Fig. 5). Coumarin exhibited the most substantial stimulatory effect on FA synthesis, accompanied by the upregulation of *FUB3*, *FUB6*, and *FUB9*. These findings suggest that coumarin may enhance the activity of amino acid kinase and NADH-dependent dehydrogenase in the FA biosynthetic pathway^[36]^. In treatments with ferulic acid, significant upregulation of *FUB1* and *FUB9* was observed, indicating that ferulic acid may augment the activity of polyketide synthase. Conversely, FA production was significantly diminished under 7-Hydroxycoumarin treatments, likely due to the inhibition of *FUB6* expression, suggesting that 7-Hydroxycoumarin may mitigate the phytotoxicity of *Foz*.

**Fig. 5.**
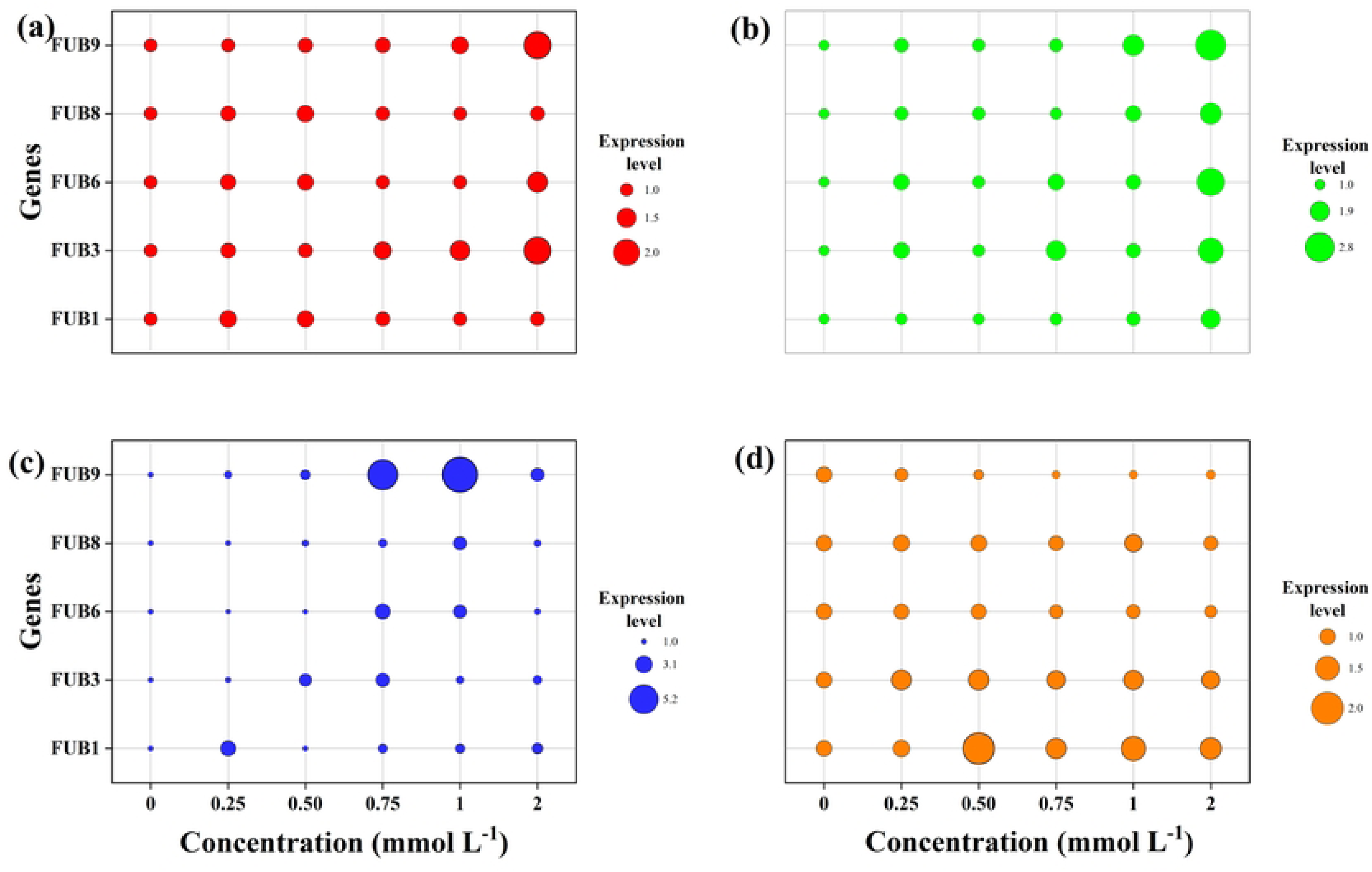
Effect of different autotoxins concentrations on the expression of *FUB* gene cluster of *Fusarium oxysporum* f. sp. *niveum (Foz).* (a): Syringic acid, (b): Coumarin, (c): Ferulic acid, (d) 7-Hydroxycoumarin.

According to the RI indices, syringic acid markedly stimulated mycelial growth and the activity of cell wall-degrading enzymes, ferulic acid significantly enhanced sporulation and biomass production, coumarin induced mycotoxin synthesis; and 7-Hydroxycoumarin facilitated conidial germination (Fig. 6). The allelopathic effects were specific to each compound, with each autotoxin exerting a distinct influence. This phenomenon may also be attributed to the different mechanisms involved in the interactions between plants and pathogens.

**Fig. 6.**
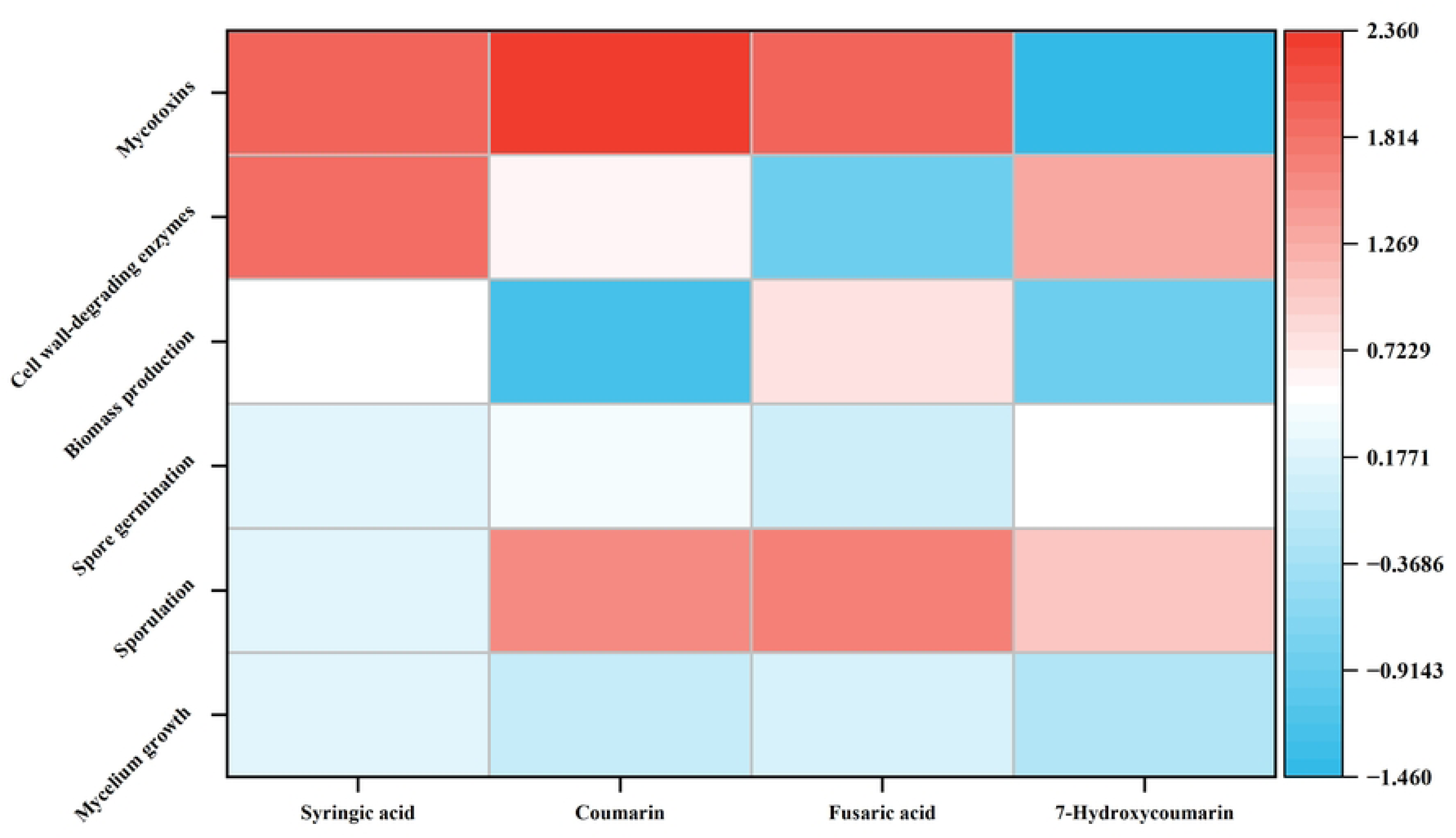
Comprehensive effect of different autotoxin on *Fusarium oxysporum* f. sp. *niveum* (*Foz*), covering mycelium growth, sporulation, conidial germination, biomass production, cell wall–degrading enzyme activity and mycotoxin synthesis.

In conclusion, the autotoxic compounds syringic acid, ferulic acid, coumarin, and 7-hydrocoumarin exert diverse allelopathic effects on the growth, reproduction, and virulence of *Fusarium oxysporum* f. sp. *zingiberi* (*Foz*), thereby exacerbating the threat of *Fusarium* wilt in ginger (Fig. 7). Notably, the allelopathic responses of *Foz* to these four compounds varied at the cellular level. Syringic acid notably enhanced mycelial growth and cell wall-degrading enzyme activity, and promoted mycotoxin production, thereby reinforcing pathogenicity. Ferulic acid and 7-hydrocoumarin significantly facilitated the growth and reproduction of *Foz*, providing ample resources for its invasion. In contrast to the other compounds, coumarin exhibited the most pronounced stimulation of mycotoxin production, potentially intensifying the symptoms of infection. Consequently, these autotoxic compounds are sufficient to promote ginger wilt disease.

**Fig. 7.**
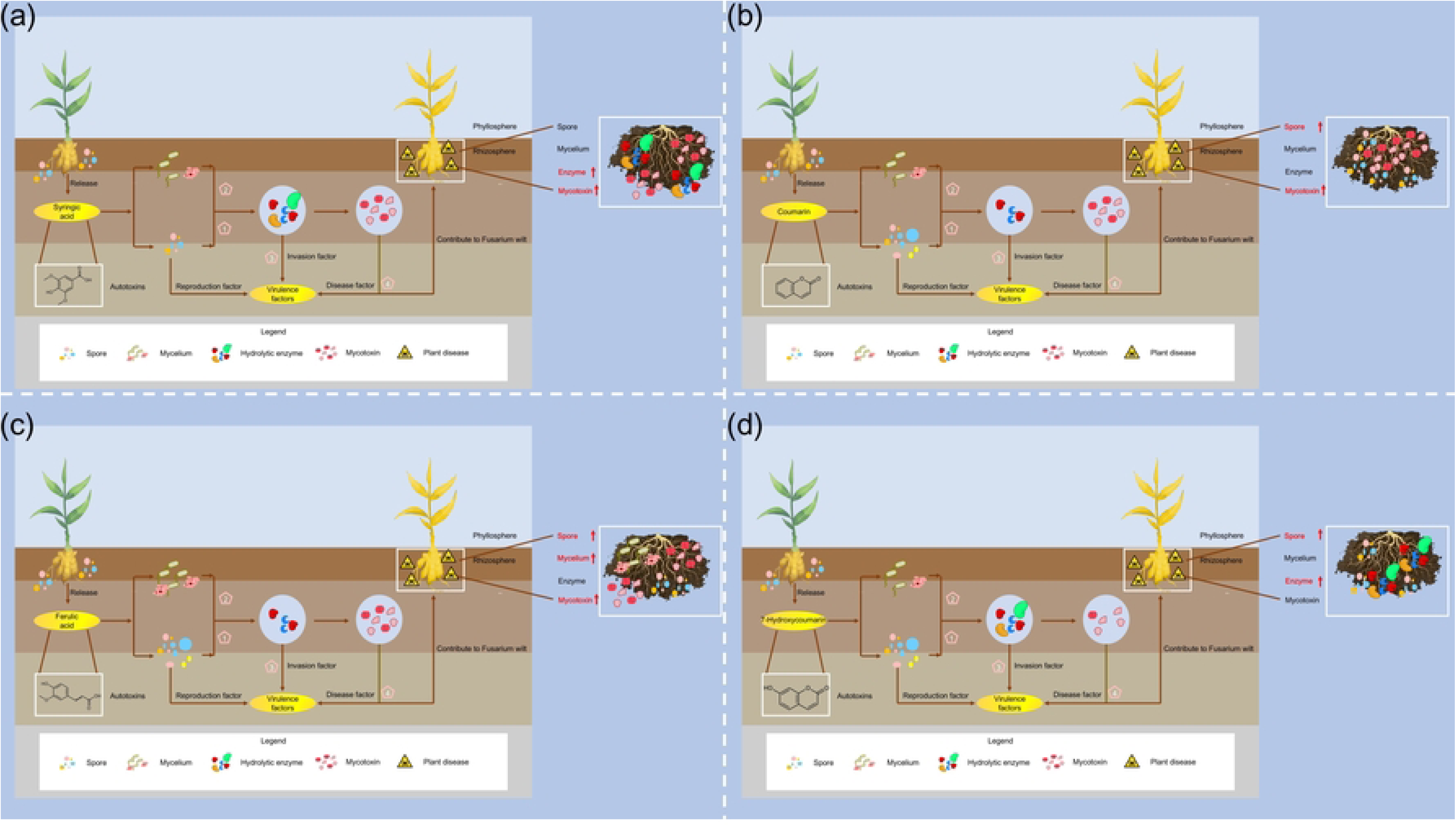
Schmatic diagram of effect of different autotoxins on *Fusarium oxysporum* f. sp. *niveum* (*Foz*). (a): Syringic acid (b): Coumarin (c): Ferulic acid; (d): 7-Hyoxycoumarin

